# Crystal Structure of INTS3/INTS6 complex reveals Functional Importance of INTS3 dimerization in DSB Repair

**DOI:** 10.1101/2020.04.10.035014

**Authors:** Yu Jia, Zixiu Cheng, Sakshibeedu R Bharath, Qiangzu Sun, Nannan Su, Jun Huang, Haiwei Song

## Abstract

SOSS1 is a single-stranded DNA (ssDNA)-binding protein complex that plays a critical role in double-strand DNA break (DSB) repair. SOSS1 consists of three subunits: INTS3, SOSSC, and hSSB1 with INTS3 serving as a scaffold to stabilize this complex. Moreover, the integrator complex subunit 6 (INTS6) participates in the DNA damage response through direct binding to INTS3 but how INTS3 interacts with INTS6 thereby, impacting DBS repair is not clear. Here, we determined the crystal structure of the C-terminus of INTS3 (INTS3c) in complex with the C-terminus of INTS6 (INTS6c) at a resolution of 2.4 Å. Structure analysis revealed that two INTS3c subunits dimerize and interact with INTS6c via conserved residues. Subsequent biochemical analyses confirmed that INTS3c forms a stable dimer and INTS3 dimerization is important for recognizing the longer ssDNA. Perturbation of INTS3c dimerization and disruption of the INTS3c/INTS6c interaction, impair the DSB repair process. Altogether, these results unravel the underappreciated role of INTS3 dimerization and the molecular basis of INTS3/INTS6 interaction in DSB repair.

## Introduction

DNA double-strand breaks (DBSs) arise when the integrity of genomic DNA is challenged by a variety of endogenous and exogenous DNA damaging agents such as replication fork collapse, oxidative stress and ionizing radiation (IR) (Aguilera and Gomez-Gonzalez, 2008). DBSs are highly cytotoxic lesions that disrupt the continuity of chromosome (van Gent et al., 2001). Defective DBS repair leads to genome instability and are associated with developmental disorders, premature aging, and tumorigenesis (Jackson and Bartek, 2009; McKinnon, 2009).

In eukaryotes, two main pathways, homologous recombination (HR) and non-homologous end joining (NHEJ) have evolved to repair DSBs (Ceccaldi et al., 2016; Chapman et al., 2012; Symington and Gautier, 2011). As a more accurate mechanism, HR requires a sister chromatid as the template to ensure genome integrity, but only occurs in the S and G2 phases of the cell cycle (Huertas, 2010). One of the initial steps in the process of HR is the resection of DSBs to generate a 3’-single-stranded DNA (ssDNA) overhang, which is essential for Rad51-mediated strand exchange (Jazayeri et al., 2008; Krejci et al., 2012; Lee and Paull, 2005). Following the resection of DSBs, the replication protein A (RPA) complex immediately binds to these ssDNA overhangs, therefore not only protecting these ssDNA regions from nucleases but also facilitating Rad51 filament formation (Zou et al., 2006). In addition to RPA, human SSB1 and SSB2 (hSSB1, hSSB2) have been shown to be involved in DNA repair mechanisms in eukaryotic cells (Richard et al., 2008). These SSBs possess an oligonucleotide/oligosaccharide binding (OB)-fold, which facilitates protein-nucleic acid, as well as protein-protein interactions. Following DNA damage, hSSB1 is stabilized by ATM-dependent phosphorylation and localizes to the sites of damaged DNA as discrete foci (Richard et al., 2008; Richard et al., 2011a; Richard et al., 2011b).

We and others have shown that hSSB1 and hSSB2 form an independent trimeric complex referred to as SOSS1 or SOSS2, with two other proteins, integrator complex subunit 3 (INTS3) and SOSSC (Huang et al., 2009; Li et al., 2009; Zhang et al., 2009). We reconstituted a SOSS1 subcomplex containing the N-terminal half of INTS3 (INTS3_N_) and full-length hSSB1, and a trimeric complex composed of INTS3_N_/hSSB1 and full length SOSSC. We determined the crystal structures of the trimeric complex (INTS3_N_/hSSB1/SOSSC) in *apo* form and in complex with a 35 nt ssDNA as well as the structure of the dimeric complex (INTS3_N_/hSSB1) complexed with a 12 nt ssDNA (Ren et al., 2014). These structures combined with functional analysis confirmed that INTS3_N_ acts as a scaffold to bridge the interaction between hSSB1 and SOSSC, and showed that hSSB1 binds to INTS3_N_ and ssDNA through two distinct surfaces (Ren et al., 2014). Recent biochemical characterization of INTS3 showed that the C-terminus of INTS3 (INTS3c) is required for nucleic acid binding (Vidhyasagar et al., 2018). However, how INTS3 contributes to ssDNA binding and the mechanism by which SOSS1 recognizes longer ssDNA remain elusive. Stringent affinity purification coupled with mass spectrometry experiments suggested yet another protein, named integrator complex subunit 6 (INTS6) to be associated with the SOSS1 (Zhang et al., 2013). In response to DSBs, INTS6 is recruited to DSBs by directly interacting with INTS3c. Depletion of INTS6 and its paralog DDX26b abolished RAD51 foci formation suggesting that INTS6 recruitment to DSBs along with INTS3 and hSSB1 regulate the accumulation of RAD51 and the subsequent HR repair (Zhang et al., 2013).

To clarify the molecular mechanism underlying the assembly of INTS3/INTS6 complex in DSB repair, we crystallized and determined the crystal structure of the INTS3c (residues 560-905, 915-995)/INTS6c (residues 800-887) complex. The structure shows that INTS3c forms a dimer and both subunits mediate its interaction with INTS6c. Biochemical assays further confirmed that INTS3 functions as a dimer. Structure-based mutagenesis indicated that mutations that disrupt the INTS3 dimer or the INTS3c/INTS6c interactions impaired RAD51 foci formation and sensitized cells to IR. Dimerization of INTS3 is also required for recognition of longer ssDNA. Overall, these findings highlight the important role of INTS3 dimer and the molecular basis of the INTS3/INTS6 interaction in DSB repair.

## Results

### Overall structure of the INTS3c/INTS6c complex

The purified C-terminal half of INTS3 (560-995) (**Fig. 1A**) was subjected to limited proteolysis followed by N-terminal sequencing to identify two stable fragments of INTS3, residues 560-905 and residues 915-995. These fragments were co-expressed and purified along with C-terminal residues 800-887 of INTS6 (**Fig. 1A**) to yield a homogenous, stable and crystallizable INTS3c/INTS6c complex. Analytical size chromatography showed that the INTS3c/INTS6c complex was eluted as a dimer of heterodimeric INTS3c/INTS6c (***SI Appendix*, Fig. S1**). Crystal structure of selenomethionine substituted INTS3c in complex with INTS6c was determined using single wavelength anomalous dispersion (SAD) method. The structure of INTS3c/INTS6c complex reveals an all α-helical assembly resembling the shape of a “butterfly”, with the INTS3c dimer forming the wings, and one INTS6c subunit near the 2-fold axis of the dimer interface making the body of the butterfly (**Fig. 1B**). INTS3c adopts a right-handed super-helical structure with 20 helices, in which each helix is paired with its neighbouring helix to form an antiparallel helix pair (**Fig. 1B**). INTS6c contains four α-helices (**Fig. 1B**) and interacts with INTS3c via two loops between α1-α2 and α3-α4.

**Fig. 1.**
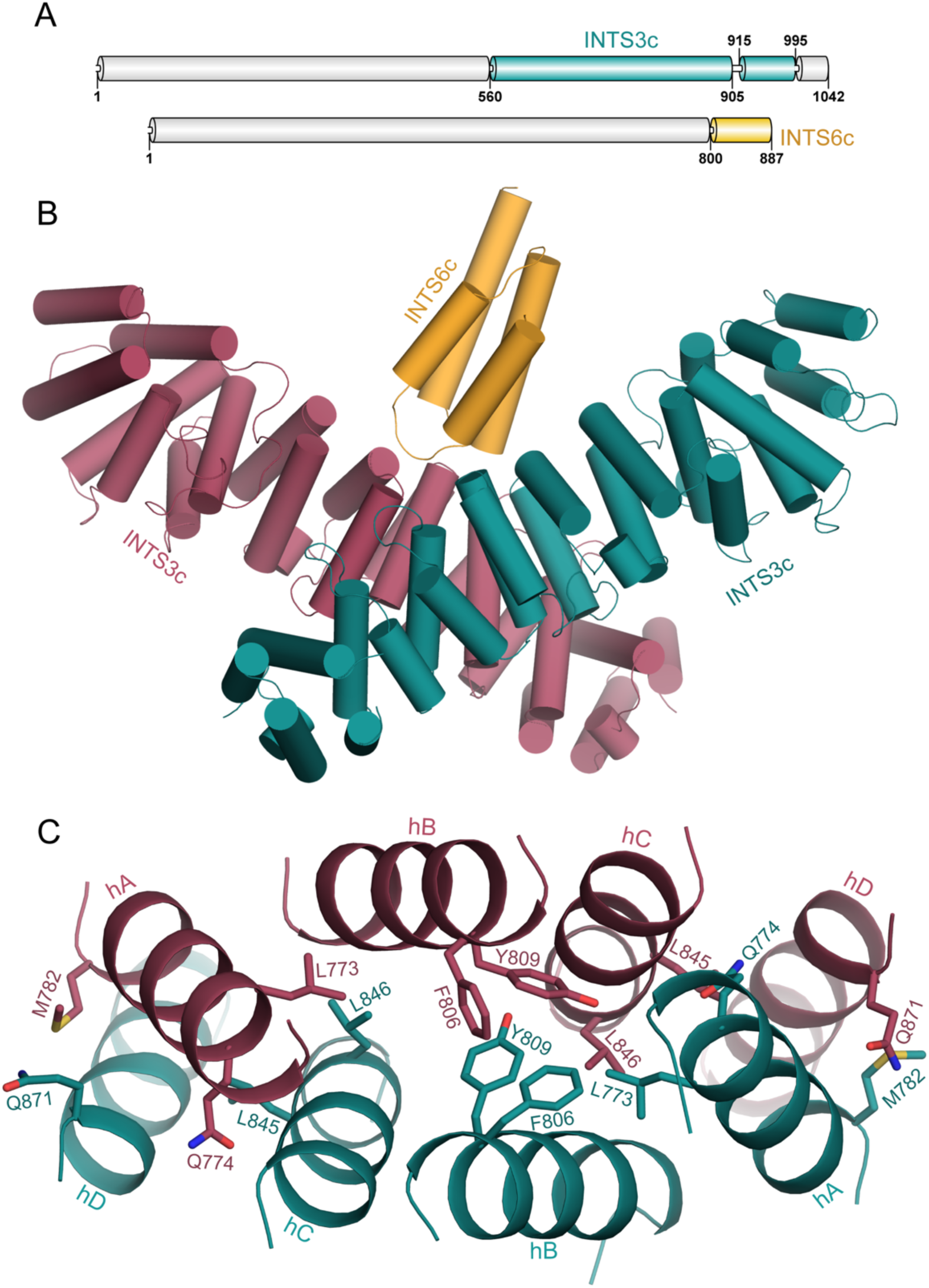
Structure of INTS3c-INTS6c. (**A**) Schematic representation of human INTS3 and INTS6. (**B**) Crystal structure of the INTS3c/INTS6c complex (**C**) The dimer interface of INTS3c. The central core of the dimeric interface of INTS3c is made up of helical segments, hA, hB, hC and hD. For simplicity of viewing, only few interactions involving Leu773, Gln774, Met782, Phe806, Tyr809, Leu845, Leu846, Gln871 and their dyad symmetry mates are shown.

### The INTS3c dimer interface

Two INTS3c protomers form a symmetrical dimer (**Fig. 1B**) wherein two protomers are related by a non-crystallographic 2-fold axis. Analysis of the dimer interface using PISA (Krissinel and Henrick, 2007) revealed a buried surface area of ∼3400.0 Å^2^. Each protomer contributes 46 residues to the dimer interface, forming a vast network of hydrophobic, van der waals, and hydrogen bond interactions with a free energy of dissociation, ΔG^diss^ of 9.7 kcal/mol. The interface mainly consists of residues belonging to four helical segments **(Fig. 1C)**, hA (Asp769-Leu785), hB (Glu804-Ile820), hC (Lys833-Lys852) and hD (Pro868-His884). The bulk of the hydrophobic core consists of Leu773, Phe806, Tyr809, and Leu846 from both chains of the dimer **(Fig. 1C)**. These interactions are further strengthened by additional hydrophobic interactions, 22 hydrogen bonds and numerous van der Waals interactions, e.g. Met782 of hA contacts Gln871 of hD and Gln774 of hA contacts Leu845 of hC. The tight INTS3c dimer and the conserved residues in the interface (***SI Appendix*, Fig. 2A**) suggest that the dimer likely corresponds to a physiologically relevant form of INTS3.

**Fig. 2.**
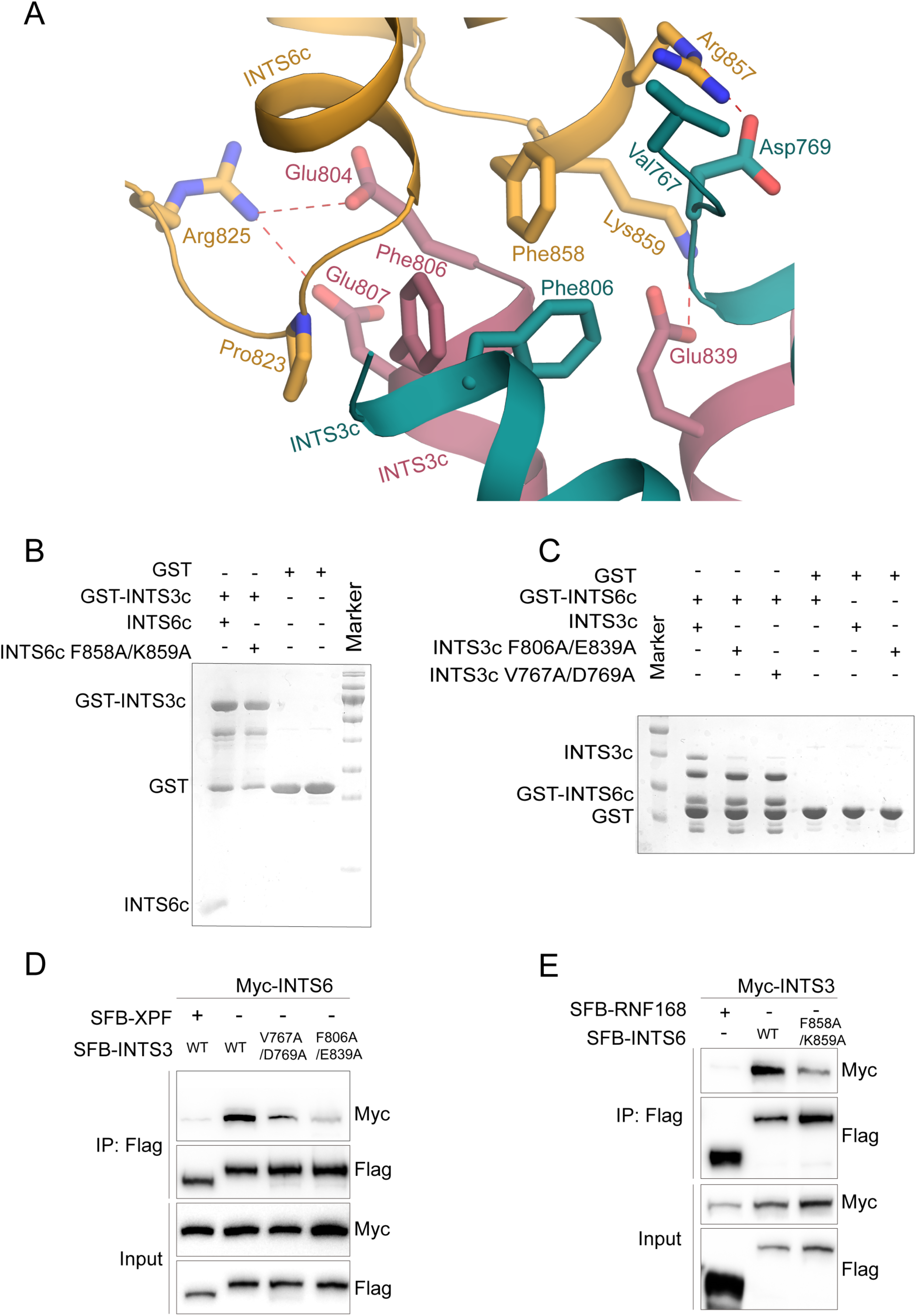
Interface between INTS3c and INTS6c. (**A**) The INTS3c-INTS6c interface is composed of a central hydrophobic residues, Pro823, Phe858 from INTS6c and Phe806 from INTS3c. These hydrophobic interactions are further strengthened by salt-bridges and hydrogen bonds. (**B**) GST-pull down assays show that the F858A/K839A mutant of INTS6c did not interact with INTS3c (**C**) Results of GST-pull down assays show that the V767A/D769A and F806A/E839A mutants of INTS3c did not interact with INTS6c. (**D**) The V767A/D769A and F806A/E839A mutants of full length INTS3 show weak interaction with full length INTS6 in HEK293T cells that were transfected with plasmids encoding Myc-tagged INTS6 and SFB-tagged wild-type INTS3 (WT) or its mutants. SFB-tagged XPF was used as a negative control. Cell lysates were immunoprecipitated with anti-Flag antibody and western blot analysis was performed with anti-Flag and anti-Myc antibodies. (**E**) The F858A/K859A mutant of full length INTS6 shows weak interaction with full length INTS3. Co-immunoprecipitation experiments were performed similar to those described in (**D**). SFB-tagged RNF168 was used as a negative control.

### The INTS3-INTS6 interface

The interactions of INTS6c with chain A and B of INTS3c account for a total buried surface area of 706.8 Å^2^ and 503.5 Å^2^, respectively. The interface is rather small with only the loops between the helical segments of INTS6c interacting with the INTS3c dimer. As shown in **Fig. 2A**, the core of the interface is hydrophobic in nature with Pro823 and Phe858 of INTS6c juxtaposed against Phe806 from the two subunits of INTS3c. The interaction is further augmented by salt bridges between Lys859 of INTS6c with Glu839 of INTS3c; Arg825 of INTS6c with Glu804 and Glu807 of INTS3c; and Arg857 of INTS6c with Asp769 of INTS3c. Sequence alignment of INTS6c from human, mouse, zebrafish and drosophila species **(*SI Appendix*, Fig. S2B)** highlighted the conserved nature of the residues involved in the interaction with INTS3c.

### Mutational Analysis

To identify the residues important for INTS3c-INTS6c interaction, we prepared a number of site-specific mutations in either INTS3c or INTS6c and examined the binding of these mutants to their respective wild-type binding partner using GST-pulldown assays. Substitution of amino acids Phe858 and Lys859 to Ala (F858A/K859A) in INTS6c abolished its binding to INTS3c **(Fig. 2B)**. Conversely, substitution of amino acids F806 and E839 to Ala (F806A/E839A) or substitution of amino acidsV767 and D769 to Ala (V767A/D769A) in INTS3c disrupted its interaction with INTS6c **(Fig. 2C)**.

We further introduced these mutations into full length INTS3 and INTS6 and tested their interaction by co-immunoprecipitation assays. As shown in **Fig. 2D**, the V767A/D769A or F806A/E839A mutant of INTS3 failed to interact with INTS6. Similarly, the F858A/K859A mutant of INTS6 also exhibited reduced binding to INTS3 **(Fig. 2E)**. Altogether, these results indicate that these residues are indeed important for the INTS3-INTS6 complex formation.

Next, we examined whether the INTS3-INTS6 interaction are important for their recruitment to DNA damage sites. As shown in **Figs. 3A-3B**, similar to wild-type INTS3, both the V767A/D769A and F806A/E839A mutants could still relocate to DSB sites following IR treatment. In contrast, the F858A/K859A mutant of INTS6 showed a diffuse cytoplasmic localization, suggesting that forming a complex with INTS3 is essential for INTS6 nuclear localization (**Fig. 3C**). These results are consistent with the previous finding that INTS3 recruits INTS6 to DSB sites in response to IR (Zhang et al., 2013). We further tested whether INTS3 damage recruitment is dependent on its dimerization. To this end, we generated two deletion mutants rather than site-directed mutants due to the extensive interface of the INTS3 dimer. Specifically, we separately deleted residues 791-831 and residues 832-872 (designated as INTS3^Δ791-831^and INTS3^Δ832-872^), which contain helices hB and hC critical for INTS3 dimerization (**Fig. 1C**). As shown in **Figs. 3A-3B**, both the deletion mutants of INTS3 failed to form IR-induced foci, indicating that dimerization is necessary for INTS3 recruitment to DSB sites.

**Fig. 3.**
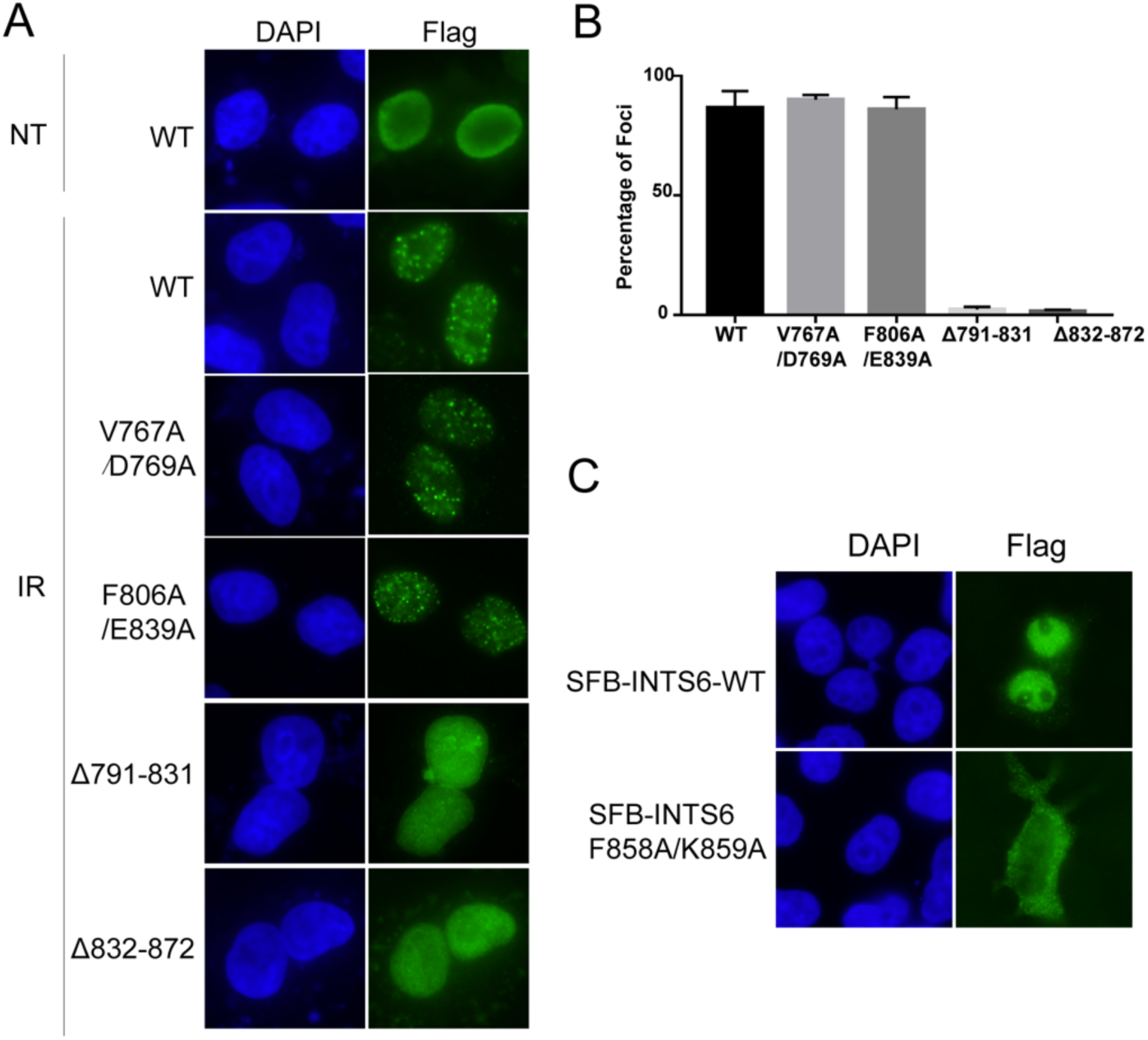
The effects of INTS3 dimerization and its interaction with INTS6 on INTS3/INTS6 recruitment to DSB sites. (**A-B**) Cells transfected with SFB-tagged wild-type INTS3 or its mutants were treated with IR (10 Gy) for 6 h or left untreated. Cells were then fixed and processed for immunofluorescence with anti-Flag antibody (A). The quantification of foci-positive cells was performed by counting a total of 300 cells per sample (B). (**C**) Cells transfected with SFB-tagged wild-type INTS6 or the F858A/K859A mutant were fixed and processed for immunofluorescence with anti-Flag antibody.

To investigate the biological significance of INTS3 dimerization and its interaction with INTS6, we utilized a doxycycline-inducible expression system to express the siRNA-resistant wild-type INTS3, INTS6 or the mutants defective in INTS3 dimerization or INTS3-INTS6 interaction in INTS3- or INTS6-depleted cells. As shown in **Figs. 4A-4B**, neither the V767A/D769A and F806A/E839A mutants nor the INTS3^Δ791-831^and INTS3^Δ832-872^ mutants were able to restore RAD51 foci formation in INTS3-depelted cells. Likewise, RAD51 foci formation in INTS6-depleted cells could be restored by wild-type INTS6 but not the F858A/K859A mutant (**Figs. 4C, 4D**). More importantly, perturbation of INTS3 dimerization or disruption of the INTS3-INTS6 complex formation rendered cells more sensitive to IR (**Figs. 4E-4F**). Taken together, these results suggest that both INTS3 dimerization and the INTS3-INTS6 complex formation are important for DSB repair.

**Fig. 4.**
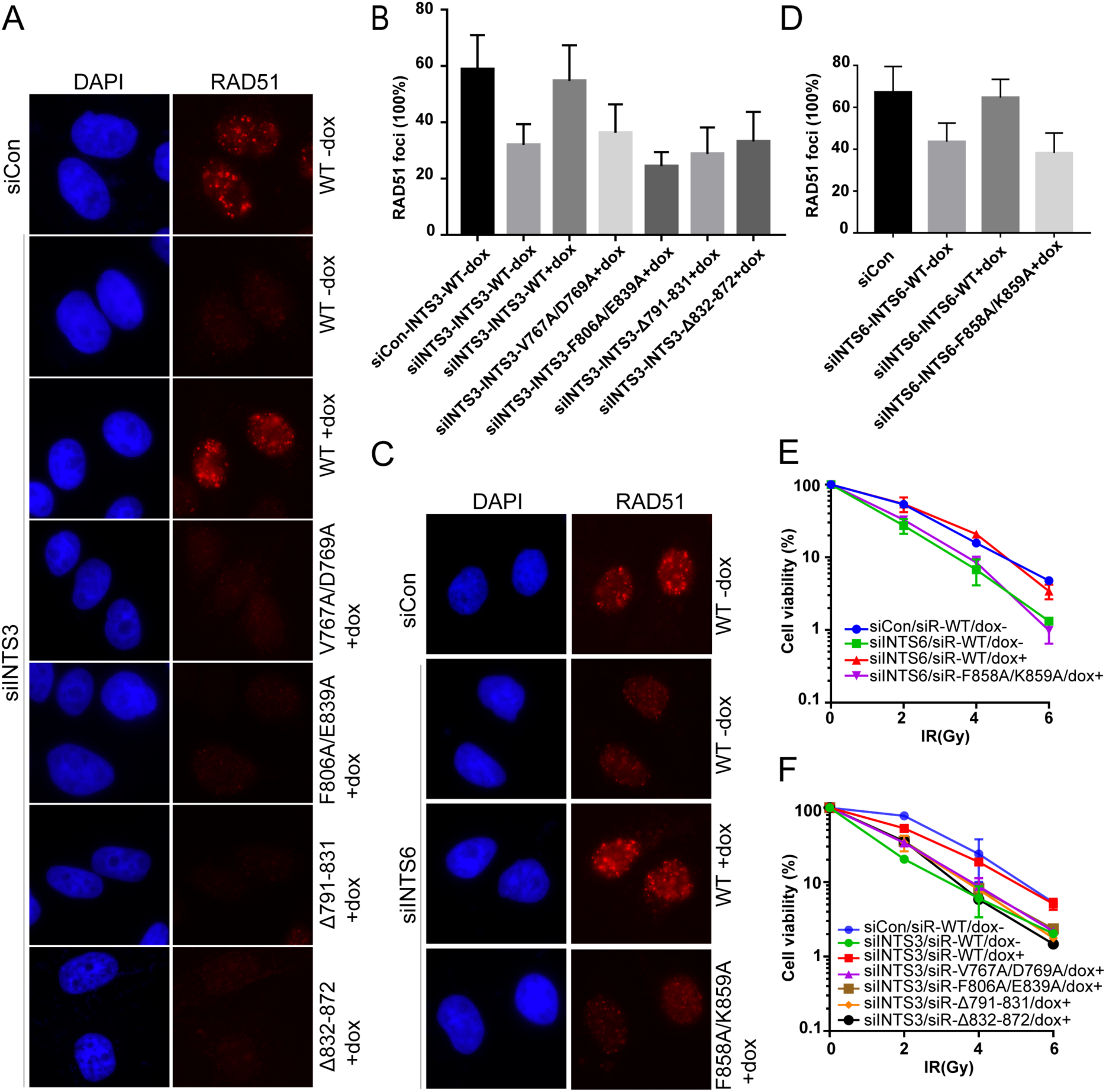
INTS3 dimerization and its interaction with INTS6 are required for DNA repair. (**A-B**) Cells stably expressing siRNA-resistant wild-type INTS3 or its mutants defective in INTS6 binding (V767A/D769A, F806A/E839A) or dimer formation (Δ791-831, Δ832-872) under the control of a tetracycline-inducible promoter were generated. The cell lines were transfected twice with INTS3 siRNA. 24 h after the second transfection, cells were cultured with doxycycline for 24 h and treated with IR (10 Gy) for 6 h. Cells were then fixed and processed for RAD51 immunofluorescence (A). The quantification of foci-positive cells was performed by counting a total of 300 cells per sample (B). (**C-D**) Cells that express siRNA-resistant wild-type INTS6 or its point mutant defective in binding to INTS3 (F858A/E859A) under the control of a tetracycline-inducible promoter were generated. The resulting cell lines were transfected twice with INTS6 siRNA. 24 h after the second transfection, cells were cultured with doxycycline for 24 h and treated with IR (10 Gy) for 6 h. Cells were then fixed and processed for RAD51 immunofluorescence (C). The quantification of foci-positive cells was performed by counting a total of 300 cells per sample (D). (**E-F**) Cells that express siRNA-resistant wild-type INTS3/INTS6 or their mutants under the control of a tetracycline-inducible promoter were generated. The resulting cell lines were transfected twice with control or INTS3/INTS6 siRNAs. After IR treatment, cells were incubated for 14 days before staining. Experiments were done in triplicates. Results shown are averages of three independent experiments.

### Recognition of ssDNA

Our previous structural studies of the SOSS1 complex (Ren et al., 2014) did not include the C-terminal domain of INTS3 resulting in poor understanding of long ssDNA recognition. Moreover, the C-terminal of INTS3 has been reported to be required for ssDNA binding (Vidhyasagar et al., 2018). To explore the function of INTS3 dimer in recognition of longer ssDNA, we created a series of deletion mutants of INTS3 to test if its dimerization could improve longer ssDNA binding by electrophoretic mobility shift assay (EMSA).

Wild-type hSSB1, INTS3(1-995)-hSSB1 and INTS3(1-500)-hSSB1 were co-expressed and purified **(*SI Appendix*, Fig. 3A)** and subjected to analytical size exclusion chromatography (SEC) analysis. INTS3(1-995)-hSSB1 contains residues of 1-995 (deletion of the C-terminal unstructured region; residues 996-1042) and a full-length hSSB1 while INTS3(1-500)-hSSB1consists of just INTS3_N_ and a full-length hSSB1 and has been shown to be in a heterodimeric form in solution previously (Ren et al 2016). As expected, INTS3(1-995)-hSSB1 forms a dimer of heterodimer while INTS3(1-500)/hSSB1 is a heterodimer in solution **(*SI Appendix*, Fig. S1)**. To examine whether dimerization affects the binding of longer ssDNA, we mixed INTS3(1-995)-hSSB1, INTS3(1-500)-hSSB1, hSSB1 with either dT12 or dT48 at different molar ratios and ran EMSA gels. INTS3(1-995)-hSSB1 binds to dT48 with the strongest affinity in comparison with INTS3(1-500)-hSSB1 or hSSB1 **(*SI Appendix*, Fig. S3C)** while INTS3(1-995)-hSSB1, INTS3(1-500)-hSSB1 and hSSB1 bind to dT12 with comparable affinities (***SI Appendix*, Fig. S3D**). Moreover,the presence of INTS6c had no effect on the binding of INTS3(1-995)-hSSB1 to dT48 (***SI Appendix*, Fig. S3C)**, suggesting that INTS6c is not required for ssDNA binding. Consistent with the EMSA results, SPR assays showed that hSSB1 and INTS3(1-500)/hSSB1 bind to dT48 with comparable affinities while INTS3(1-995)-hSSB1 and full length INTS3/hSSB1 bind to dT48 with ∼20 times or 4 times stronger affinities than that of hSSB1 alone or INTS3(1-500)/hSSB1, respectively (**Fig. 5**). To further clarify the role of INTS3 dimerization in the binding of longer ssDNA, we co-expressed INTS3^Δ791-831^and INTS3^Δ832-872^ in the context of a truncated INTS3 (residues 1-995) with hSSB1 (termed as INTS3 (1-995)^Δ791-831^-hSSB1 and INTS3 (1-995)^Δ832-872^-hSSB1 respectively). Purified INTS3 (1-995)^Δ791-831^-hSSB1 and INTS3 (1-995)^Δ832-872^-hSSB1 showed similar patterns to that of wild-type INTS3 (1-995)-hSSB1, suggesting that the deletions in the INTS3 dimer interface did not affect protein folding so much (***SI Appendix*, Fig. S3B**). EMSA experiments showed that both INTS3 (1-995) ^Δ791-831^-hSSB1 and INTS3 (1-995) ^Δ832-872^-hSSB1 exhibited dramatically reduced binding to dT48 compared to INTS3(1-995)-hSSB1 **(*SI Appendix*, Fig. S3B)**. Altogether, these results strongly suggest that dimerization of INTS3 is required for the binding of the longer ssDNA but has no effect on the short ssDNA binding.

**Fig. 5.**
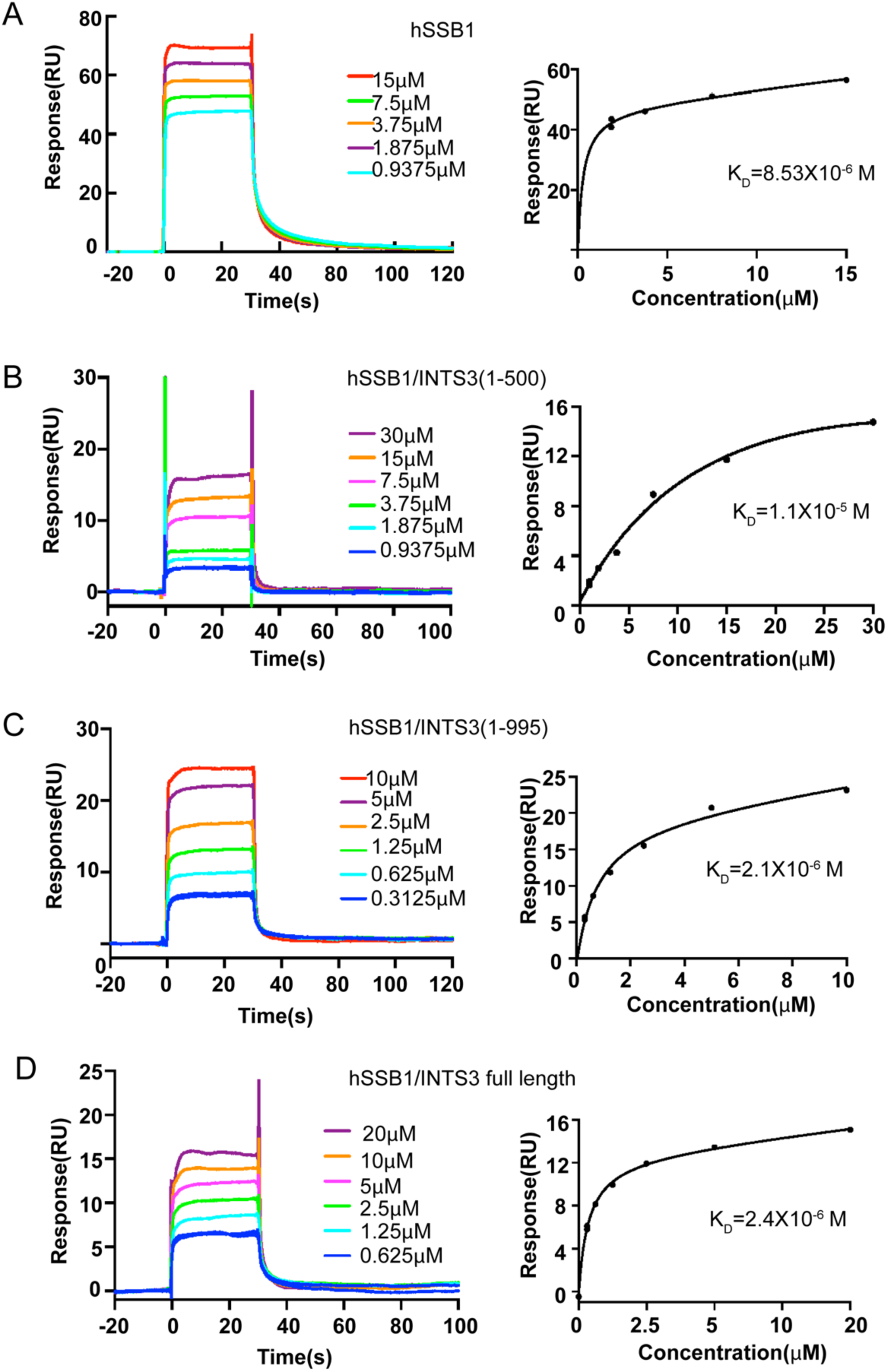
Interaction of hSSB1, INTS3/hSSB1 and its mutants with ssDNA. Sensograms resulting from Surface Plasmon resonance (SPR) assay for interaction between (**A**) immobilized hSSB1 and dT48 (B) immobilized INTS3(1-500) and dT48 (C) immobilized INTS3 (1-995)/hSSB1 and dT48 and (D) immobilized full length INTS3/hSSB1 and dT48. Dissociation constant (K_D_) was calculated by fitting to a one-site binding model.

### Mechanistic insights into longer ssDNA recognition by SOSS1 complex

A striking feature of the structure of INTS3c/INTS6c complex is the formation of an X-shaped INTS3 dimer. Importantly, we demonstrated the functional importance of INTS3 dimerization in DSB repair as well as in recognizing the longer ssDNA. Our previous crystal structure of INTS3_N_/SSB1/SOSSC showed that recognition of ssDNA with a length up to nine nucleotides is solely mediated by a single OB domain of hSSB1 (Ren et al., 2014). But how the full-length SOSS1 complex recognizes the longer ssDNA up to 35 nucleotides remains unclear. hSSB1 is a monomer under reducing conditions (Richard et al., 2008; Richard et al., 2009). However, oxidation stress can promote hSSB1 to form a tetramer, involved in oxidative DNA damage repair and is similar to other tetrameric SSBs (Paquet et al., 2016; Touma et al., 2017). The tetrameric SSBs from *E. coli* (*Ec*-SSB) and *P. falciparum (Pf-SSB)* consist of two distinct dimer interfaces, an extensive interface formed by subunits I and II (or subunits III and IV) with a buried surface area of 1200 Å^2^ and a significantly smaller interface formed by subunits II and III (or subunits I and IV) with a buried surface area of 600 Å^2^ (Antony et al., 2012; Raghunathan et al., 2000). The I-II SSB dimer binds to a 35-nt ssDNA in *Ec*-SSB and *Pf-SSB* (Antony et al., 2012; Raghunathan et al., 2000), suggesting that this SSB dimer is functionally important. Further support for this notion comes from the structure of *D. radiodurans* SSB wherein, a single polypeptide chain forms two OB domains-analogous to the I-II dimer in *Ec*-SSB and *Pf-*SSB (George et al., 2012).

We speculated that in the full-length SOSS1 complex, INTS3 dimerization would force hSSB1 to assemble into a dimer with its interface similar to the I-II interface in *Ec*-SSB and *Pf-SSB*. To explore the mechanism governing the longer ssDNA recognition by the SOSS1 complex, we superimposed the hSSB1 subunit of INTS3_N_/SSB1/SOSSC onto each SSB monomer in the *Pf-SSB* tetramer as the polarity of ssDNA in the structure of *Ec*-SSB is opposite to those in all other known structures of tetrameric SSBs. The results revealed that superposition of hSSB1 subunit of INTS3_N_/SSB1/SOSSC onto subunits I and II or subunits III and IV leads to a plausible model whereas other superpositions resulted in serious steric clashes. We then manually aligned the dyad axis of the hSSB1 dimer with that of the INTS3c dimer to generate the final INTS3/hSSB1/SOSSC/INTS6c model (**Fig. 6**). In support of the validity of this model, the ssDNA of ∼35 nt modelled to hSSB1 contacts the residues of INTS3_N_ with the same polarity, which were previously observed in the structure of INTS3_N_/SSB1/SOSSC (Ren et al., 2014). Such a model could explain how SOSS1 binds to the longer ssDNA and at the same time allow hSSB1 to interact with the MRN complex via its C-terminal domain (Richard et al., 2011a; Richard et al., 2011b). However, this model is not compatible with the disulphide-mediated hSSB1 tetramer assembled under oxidized conditions (Touma et al., 2017). In line with this notion, tetramerization of hSSB1 has been shown to be dispensable for DSB repair (Paquet et al., 2016).

**Fig. 6.**
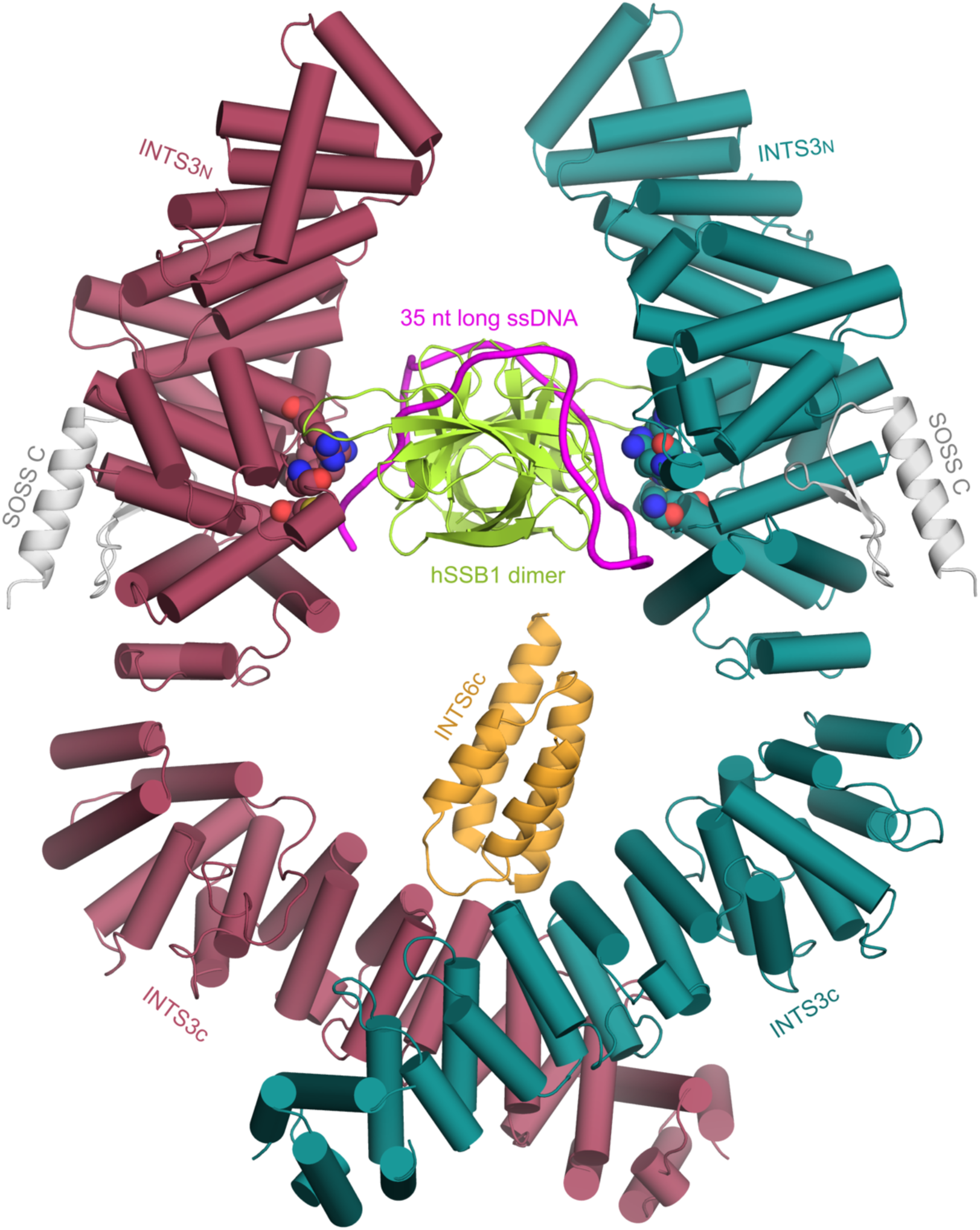
The model of INTS3/hSSB1/SOSSC/INTS6c interacting with 35 nt long ssDNA. The color coding for INTS3 and INTS6c is the same as in Figure 1. SOSSC and SSB1 are shown in light grey and limon respectively while ssDNA is shown in magenta. The residues that were previously observed to contact ssDNA in the structure of INTS3_N_/SSB1/SOSSC are shown in space filling models.

## Discussion

Our previous study showed that INTS3 is a scaffold protein with its N-terminal half mediating the interaction with hSSB1 (Ren et al., 2014). In this study, we showed that INTS3 uses its C-terminal region to mediate its interaction with INTS6. Consistent with the previous observations that the INTS3/INTS6 interaction is required for the recruitment of INTS6 to DNA damage sites in response to IR (Zhang et al., 2013), our structure-based functional assays showed that disruption of the INTS3c/INTS6c interaction inhibits the DSB repair process. Both INTS3 and INTS6 are components in Integrator complex participating in 3’ non-coding RNA processing (Baillat et al., 2005). Our studies showed that INTS6 interacts with INTS3 as a component of integrator complex involved in DSB repair. In transcription process, R-loop formed of RNA/DNA hybrids may cause transcription pause and termination and result in genome instability (Huertas and Aguilera, 2003). R-loop also accumulates at DSB site (Ohle et al., 2016). INTS6 contains a DEAD (Asp-Glu-Ala-Asp) motif usually found in RNA/DNA helicase. A Von Willebrand factor type A domain is located at the N-terminal region of INTS6, which may also serves as a platform for protein-protein interaction (Springer, 2006). This indicates that INTS3/INTS6/hSSB1 complex may have a role in binding and unwinding R-loop by recruiting other integrator members. Recently, INTS7 was found to interact with INTS3/hSSB1 as a whole complex in response to UV laser and its depletion result in cell cycle arrest bypass and mitomycin C sensitivity (Cotta-Ramusino et al., 2011). More studies are required to clarify the role of INTS3/INTS6/hSSB1 complex in DNA damage repair process.

Vidhyasagar et al showed recently that purified full-length human INTS3 is a monomer in solution (Vidhyasagar et al., 2018). In contrast, our crystal structure of INTS3c/INTS6c complex indicated that the C-terminal region of INTS3 dimerizes and such a dimerization is independent of its interaction with INTS6c. Importantly, we showed that the two interface deletion mutants of INTS3 were observed to be diffused in the cytoplasm and failed to rescue the formation of RAD51 foci, suggesting that INTS3 dimerization is critical for DSB repair.

The SOSS1 complex binds to ssDNA with a minimal length of ∼35 nucleotides (nt) and its DNA binding affinity is ∼30-fold stronger than that of hSSB1 alone (Yang et al., 2013). Structural studies of INTS3_N_/hSSB1/SOSSC (Ren et al., 2014) showed that hSSB1 solely contributes to the binding of ssDNA while neither SOSSC nor INTS3_N_ are critical for ssDNA binding, implying that the C-terminal half of INTS3 may be involved in the recognition of longer ssDNA. Consistent with this view, Vidhyasagar et al showed that the C-terminus of INTS3 rather than INTS3_N_ binds to dT30 albeit with weak affinity probably contributed by the positive charged residues at its extreme C-terminal region (Vidhyasagar et al., 2018). However, our SPR data showing that INTS3(1-995)/hSSB1 and full length INTS3/hSSB1 bind to dT48 (**Fig. 5**) with comparable affinities and purified INTS3c alone is unable to bind ssDNA (data not shown), suggesting that the C-terminus of INTS3 has no direct contribution to ssDNA binding. Instead, our EMSA and SPR results strongly indicated that the C-terminus of INTS3 is required for the binding of INTS3/hSSB1 to dT48 via dimerization. As depicted in **Fig. 6**, one simple explanation for enhanced longer ssDNA binding is that the dimerization of INTS3 engages two hSSB1 molecules to form a hSSB1 dimer with larger ssDNA recognition surface/channel for binding longer ssDNA. However, such a hSSB1 dimer is not compatible with that deposited in PDB database (no accompanying paper has been published), which shows that hSSB1 dimerizes via its C-terminal tail. Furthermore, superposition of INTS3_N_/SSB1/SOSSC with hSSB1 (PDB codes: 5D8E and 5D8F) at SSB1 give rise to severe steric clashes, suggesting that the hSSB1 dimer in the deposited structures might not be the physiologically relevant form.

Previously, it was reported that both N and C termini of INTS3 are important for its foci formation after DNA damage but its C-terminal deletion exhibited stronger phenotype than its N-terminal deletion (Zhang et al., 2009). Consistent with these results, our previous and present data show that the N-terminus of INTS3 functions as a scaffold for SOSS1 assembly while the C-terminus of INTS3 dimerizes to promote the formation of larger SOSS1 complex. The large and conserved INTS3 dimer interface plus its functional importance suggests that the dimer of INTS3 is likely to be its functional unit in DSB repair and is required for the assembly of larger SOSS1 complex. Consistent with this notion, SOSS1 exists as more than one copy *in vivo* (Li et al., 2009). The larger SOSS1 complex formation mediated by INTS3 dimerization not only is required for foci formation but also is important for recognizing longer ssDNA during DSB repair process. Further structural and functional studies on the full length INTS3 in complex with hSSB1 are required to clarify the mechanism governing the role of SOSS1 in DSB repair.

## Methods

### Purification, crystallization and structure determination of INTS3c/INTS6c

The C-terminal half of INTS3 consisting of residues 560-995, was cloned into pGEX-6p-1 vector, expressed in *E. coli* BL21 codonPlus cells and purified to homogeneity using successive chromatographic steps involving GST affinity and gel filtration columns. However, the purified C-terminal of INTS3 was prone to degradation. In an effort to characterize the most stable fragment of INTS3, we used limited proteolysis followed by N-terminal sequencing to identify two stable fragments of INTS3, residues 560-905 and 915-995. These two fragments were co-expressed in pETDuet1 vector with GST tag and the complex (INTS3c) was purified to > 95% purity. The INTS6 fragment consisting of residues 800-887 (INTS6c) was cloned into pGEX-6p-1 vector, expressed in BL21 (DE3) codonPlus cells and was purified using successive chromatographic steps involving GST affinity and gel filtration columns. The purified INTS3c and INTS6c were mixed together and the resultant INTS3c/INTS6c complex was subjected to size exclusion chromatography. Crystallization trials yielded clusters of small parallelepipeds that diffracted to ∼3 Å. However, the sequences of INTS3c and INTS6c have no known homologs in the Protein Data Bank. Hence, we incorporated selenomethionine into INTS3c using feedback inhibition of methionine biosynthesis. The selenomethionine incorporated crystals of INTS3c/INTS6c diffracted to 2.6 Å resolution. A diffraction dataset was collected at the Australian synchrotron; and dataset was processed using *XDS* and *Aimless* of CCP4. Selenomethionine sites were identified using *HySS* using single anomalous dispersion method. The sites were refined using *Resolve* and phases were extended to the whole complex using *PHASER* in PHENIX suite of crystallographic programs. The resulting structure was used as molecular replacement model for data collected on native INTS3c/INTS6c crystals that diffracted to 2.4 Å. The structure was further manually built in *COOT* and refined using *phenix.refine* to R_work_ and R_free_ values of 0.21 and 0.26, respectively. Statistics of the structure determination and refinement are summarized in ***SI Appendix*, Table S1.**

### Cell culture and transfection

HEK293T and HeLa Cells were maintained in DMEM supplemented with 10% fetal bovine serum and 1% penicillin and streptomycin. Cell transfection was performed using Lipofectamine2000 (Invitrogen), following the manufacturer’s protocol. The mammalian expression plasmids for SFB- or Myc-tagged INTS3 and INTS6 were previously described (Huang et al., 2009). Site-directed mutagenesis was performed to obtain the INTS3, INTS6 or INTS3 mutants. For transient expression of INTS3, INTS6 or their mutants, the corresponding fragment in the entry vector was transferred into Gateway compatible destination vector which harbor either an N-terminal triple-epitope tag (S protein tag, Flag epitope tag and Streptavidin binding peptide tag) or an N-terminal Myc tag. All clones were sequenced to verify desired mutations.

### RNA interference

All siRNAs were synthesized by Dharmacon Inc. The siRNAs were 21 base pairs and sequences are as follows: INTS3 siRNA: 5’-GAUGAGAGUUGCUAUGACAdTdT-3’; INTS6 siRNA: 5’-GGAAAGAAAUUGAUGCAUUdTdT-3’ and control siRNA: 5’-UUCAAUAAAUUCUUGAGGUUU-3’. The siRNA-resistant wild-type and mutant INTS3 constructs were generated by changing nine nucleotides in the INTS3 siRNA targeting region (G1569A, T1572C, G1575A, A1576T, G1577C, T1578A, C1581T, T1584C and C1587T substitutions). The siRNA-resistant wild-type and mutant INTS6 constructs were generated by changing six nucleotides in the INTS6 siRNA targeting region (A2388T, G2391A, A2394G, G2397A, T2403C and C2406G substitutions).The siRNA transfection was performed with 100 nM siRNA duplexes using LipofectamineRNAiMAX following the manufacturer’s instruction. Transfection was repeated twice with an interval of 24 hr to achieve maximal RNAi effect.

### Antibodies

Specific antibodies recognizing RAD51 are previously described (Huang et al., 2009; Wan et al., 2013). Anti-Myc (9E10) and anti-Flag (M2) antibodies were purchased from Covance and Sigma, respectively.

### Co-immunoprecipitation and western blotting

For Flag immunoprecipitations, 0.8-ml aliquot of lysate was incubated with 1 µg of the Flag monoclonal antibody and 25 µl of a 1:1 slurry of Protein A Sepharose for 2 h at 4 °C. The Sepharose beads were washed three times with NTEN buffer, boiled in 2X SDS loading buffer, and resolved on SDS-PAGE. Membranes were blocked with 5% milk in TBST buffer and then probed with antibodies as indicated.

### Cell survival assays

HeLa cells were transfected twice with control siRNA or siRNAs specifically targeting INTS3 or INTS6. 24 hours after the second transfection, cells (10^3^) were split and transferred into 60 mm dishes. After incubation for 24 h, cells were treated with IR as indicated. 24 h later, the medium was replaced and cells were incubated for further 14 days. Resulting cells were fixed and stained with Coomassie blue.

### Lentivirus packaging and Infection

Tet-On inducible SFB-tagged lentiviral vector and packaging plasmids (pMD2G and pSPAX2) were kindly provided by Prof. Songyang Zhou (Baylor College of Medicine). INTS3 entry constructs were transferred into the Gateway-compatible SFB-tagged lentiviral vector. Virus supernatant was collected 48 h after co-transfection of lentiviral vectors and packaging plasmids (pMD2G and pSPAX2) into HEK293T cells. Cells were infected with viral supernatants with the addition of 8 µg ml^-1^ polybrene (Sigma), and stable pools were selected with medium containing 500 µg ml^-1^ G418 (Calbiochem). The expression of the indicated genes in the stable pools was induced by the addition of 1 µg ml^-1^ doxycycline (Sigma) for 24 h.

### Immunofluorescence staining

Indirect immunofluorescence was carried out as described (Huang et al., 2009). HeLa cells cultured on coverslips were treated with IR (10 Gy) for 6 h. Cells were then washed with PBS, pre-extracted with buffer containing 0.5% Triton X-100 for 5 min, and fixed with 3% paraformaldehyde for 10 min at room temperature. Cells were incubated in primary antibody for 30 min at room temperature. After three 5 min wash with PBS, secondary antibody was added and incubated at room temperature for 30 min. Cells were then stained with DAPI to visualize nuclear DNA. The coverslips were mounted onto glass slides with anti-fade solution and visualized using a Nikon ECLIPSE i80 fluorescence microscope with a Nikon Plan Fluor 603 oil objective lens.

### Electrophoretic mobility shift assay

INTS3(1-995)/hSSB1 was mixed with INTS6c at the molar ratio of 1:1 on ice for 1 h. Then INTS3(1-500)/hSSB1, INTS3(full length)/hSSB1, INTS3(1-995)/hSSB1/INTS6c, INTS3(1-995)/hSSB1 or its mutants were incubated with 0.5 fmoles of 6’-carboxy-fluorescein(6’-FAM) labelled dT48 at the molar ratio of 0.25:1, 0.5:1, 1:1 and 2:1 in the buffer (50 mM NaCl, 20 mM HEPES, 1 mM MgCl2, 0.5 mM EDTA, 0.2 mM DTT, pH 7.5) on ice for 1 h. The reactions were resolved on 4-20% gradient native gel with 0.5X TBE buffer for 4 h at 110V. The gels were analyzed by Typhoon 8600.

### Surface plasmon resonance (SPR)

SPR measurements were performed at 25 °C using BIACORE T200 instrument and the data were analysed using Graphpad Prism 7.0. INTS3(1-500)/hSSB1, INTS3(1-995) /hSSB1, INTS3(full length) /hSSB1 complex were immobilized on to the Series S NTA sensor chip in the running buffer (200 mM NaCl, 40 mM HEPES, pH 7.5, 50 µM EDTA). The manual injection process lasted for 30 s at 20 ul min^-1^ to reach immobilization level of 1000RU. dT48 was dissolved in the running buffer in multiple proportion dilution. The contact time was 30 s for each concentration of dT48 and the disassociation time was 300 s at 30 µl min^-1^.

### Modeling the plausible interaction of SOSS1 complex with longer ssDNA

Human SSB1 of INTS3_N_/hSSB1/SOSSC/ssDNA (PDB ID: 4OWW) was superposed on subunits I and II of *P. falciparum* SSB (PDB ID: 3ULP) to generate a dimeric hSSB1/INTS3_N_ complex. Furthermore, the two ssDNA fragments bound to subunits I and II of *Pf* SSB were manually connected by building intervening nucleotides such that ∼35 nt long ssDNA is wound around the dimeric hSSB1. The resulting dimeric hSSB1/INTS3_N_/ssDNA complex was subjected to unrestrained molecular dynamics simulations to optimize the interactions between hSSB1 and ssDNA. The final model of SOSS1 complex interacting with a 35 nt ssDNA is generated by aligning the dyad axis of dimeric hSSB1/INTS3_N_ with that of INTS3c/INTS6c complex such that the C-terminus of INTS3_N_ is close to the N-terminus of the INTS3c with appropriate contacts.

## ACKNOWLEDGEMENTS

We would like to thank the beamline scientists at MX2, Australian Synchrotron for assistance and access to synchrotron radiation facilities. This work was supported by Natural Science Foundation of China (Grant No. 31470723; H.S) and the Agency for Science, Technology and Research in Singapore (H.S.). We thank Dr. Roland Gamsajaeger at Western Sydney University for sharing tetrameric model of hSSB1.

## AUTHOR CONTRIBUTIONS

H.S. and J.H designed the research plan. Y.J., Z.C., S.R.B, Q.Z. and N.S. performed the experiments. Y.J., Z.C., S.R.B.,J.H. and H.S. analyzed the data and wrote the manuscript.

## SI Appendix

**Table S1.** Data collection and refinement statistics.

|  | Native | Semet |
| --- | --- | --- |
| **Data collection** |  |  |
| Space group | P2<sub>1</sub> | P2<sub>1</sub> |
| Cell dimensions |  |  |
| *a*, *b*, *c* (Å) | 60.57, 126.03, 84.44 | 60.87, 125.93, 84.78 |
| $\beta$ (°) | 102.12 | 102.14 |
|  |  | *Peak* |
| Wavelength | 1.00 | 0.966 |
| Resolution (Å) | 49.45 - 2.40 (2.49 - 2.40) | 43.25 - 2.60 (2.72 - 2.60) |
| *R*<sub>merge</sub> | 0.08 (0.75) | 0.09 (0.523) |
| *I* / $\sigma I$ | 4.9 (1.3) | 8.5 (2.2) |
| CC1/2 | 0.995 (0.53) | 0.993 (0.61) |
| Completeness (%) | 90.0 (92.9) | 99.3 (99.6) |
| Redundancy | 2.7 (2.7) | 3.2 (3.2) |
| **Refinement** |  |  |
| Resolution (Å) | 49.45 - 2.40 (2.49 - 2.40) |  |
| No. reflections | 41330 (2125) |  |
| *R*<sub>work</sub> / *R*<sub>free</sub> | 0.21 / 0.26 |  |
| No. atoms |  |  |
| Protein | 6733 |  |
| Water | 214 |  |
| *B*-factors |  |  |
| Protein | 31.02 |  |
| Water | 48.96 |  |
| R.m.s deviations |  |  |
| Bond lengths (Å) | 0.006 |  |
| Bond angles (°) | 1.36 |  |
| Ramachandran plot (% residues) |  |  |
| Allowed | 97.8 |  |
| Generously allowed | 1.6 |  |
| Disallowed | 0.6 |  |

Values in parentheses are for the highest-resolution shell.

## SI Appendix

### Figures

**Fig. S1.**
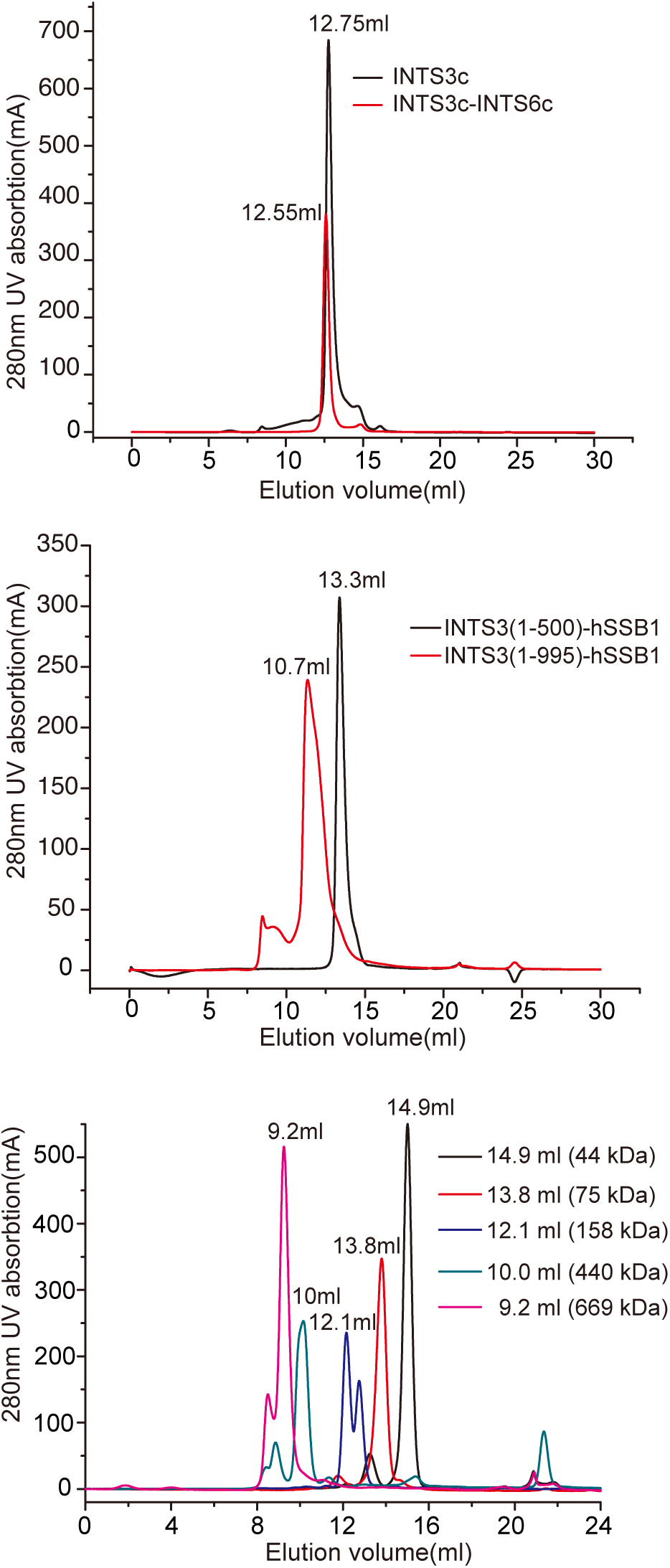
Gel filtration profiles of INTS3c, INTS3c/INTS6c, INTS3(1-500)/hSSB1 and INTS3(1-995)/hSSB1. Purified INTS3c, INTS3c/INTS6c, INTS3(1-500)/hSSB1 and INTS3(1-995)/hSSB1 elute at 12.5, 12.7, 13.3 and 10.7 ml, respectively from S200 superdex gel filtration column (top and middle panels). Comparison with elution volumes of the molecular weight markers in the same column (bottom panel) suggests that protein complexes, INTS3c, INTS3c/INTS6c and INTS3 (1-995)/hSSB1 are likely to be dimers in solution and INTS3(1-500)/hSSB1 is a monomeric complex.

**Fig. S2.**
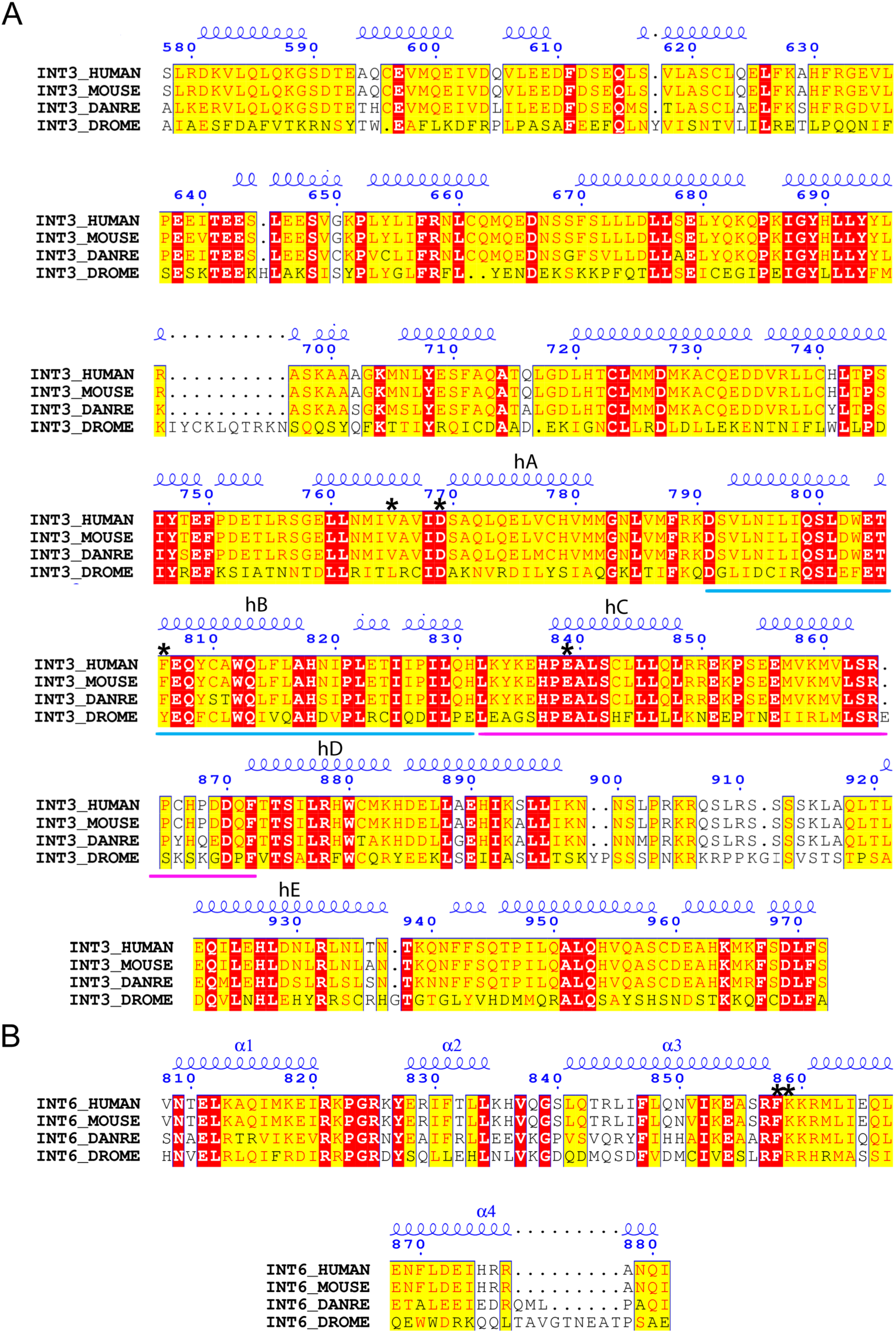
Sequence alignments of INTS3c and INTS6c with homologs from human, mouse, zebra fish and Drosophila. **(A)** Sequence alignment of human (HUMAN), mouse (MOUSE), zebrafish (DANRE) and Drosophila (DROME) INTS3c reveal conserved features. The secondary structures of human INTS3c are shown on top of the alignment. The helices involved in INTS3c dimer formation are labeled hA, hB, hC, hD and hE. **(B)** Sequence alignment of human (HUMAN), mouse (MOUSE), zebrafish (DANRE) and Drosophila (DROME) INTS6c with secondary structures of human INTS6c shown on top of the alignment. The residues mutated in the study, Val767, Asp769, Phe806, Glu839 in INTS3c and Phe858 and Lys859 in INTS6c are marked with asterisk on top of the alignment. The two segments in deletion constructs of INTS3, Δ791-831 and Δ832-872 are marked as blue and pink bars, respectively underneath the alignment.

**Fig. S3.**
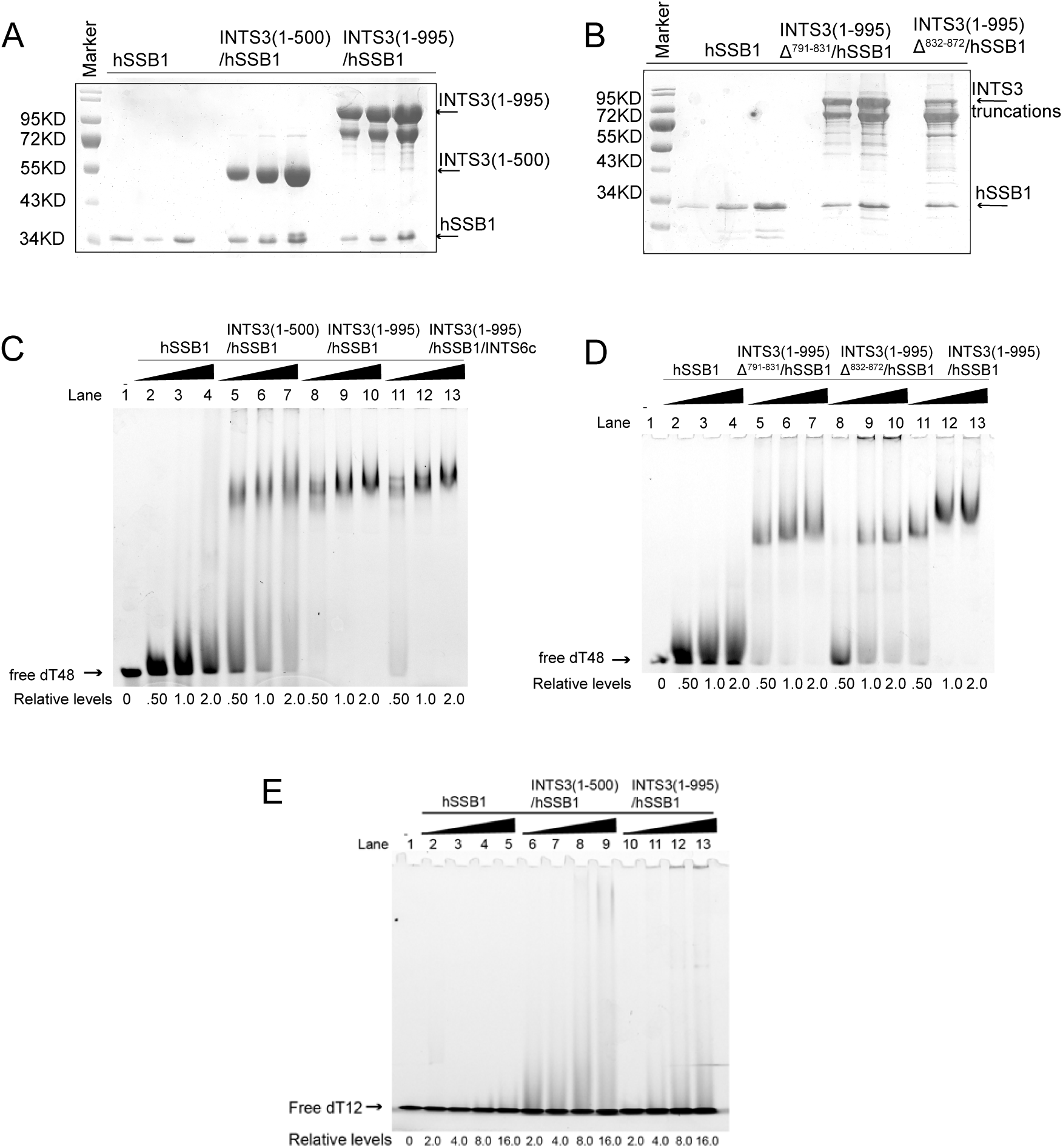
Interaction of hSSB1 in complex with different INTS3 constructs with ssDNA. **(A)** Coomassie blue stained SDS-PAGE shows hSSB1, INTS3_N_/hSSB1 and INTS3 (1-995)/hSSB1. Arrows point to protein bands. **(B)** Coomassie blue stained SDS-PAGE shows hSSB1, INTS3 (1-995) ^Δ791-831^/hSSB1 and INTS3 (1-995) ^Δ832-872^/hSSB1. Arrows point to protein bands. **(C)** Interaction of dT48 with hSSB1, INTS3_N_/hSSB1, INTS3 (1-995)/hSSB1 and INTS3 (1-995)/hSSB1/INTS6c as examined by electrophoretic mobility assay (EMSA). **(D)** Interaction of dT48 with hSSB1, INTS3 (1-995) ^Δ791-831^/hSSB1, INTS3 (1-995) ^Δ832-872^/hSSB1 and INTS3 (1-995)/hSSB1 as examined by electrophoretic mobility assay (EMSA). **(E)** Interaction of dT12 with hSSB1, INTS3_N_/hSSB1 and INTS3 (1-995)/hSSB1 as examined by electrophoretic mobility assay (EMSA). The protein samples were incubated with ssDNA on ice for 1h before loading onto the gel.

